# Transcriptomic signatures associated with mania-to-depression and depression-to-mania transitions in bipolar disorder: a case report using induced microglia-like (iMG) cells

**DOI:** 10.64898/2026.07.12.735946

**Authors:** Shogo Inamine, Sota Kyuragi, Masahiro Ohgidani, Tetsuaki Kimura, Ituro Inoue, Tomohiro Nakao, Takahiro A. Kato

## Abstract

**Introduction:** Bipolar disorder (BD) is characterized by recurring episodes of mania and depression. Despite extensive research, the pathophysiology underlying these mood swings remains elusive. Emerging evidence indicates a potential role for neuroinflammation and microglial activation in the pathophysiology of BD.

**Methods:** We employed a reverse-translational approach to generate directly induced microglia-like (iMG) cells from peripheral blood monocytes of a single patient with BD, repeatedly sampled across depressive, manic, and subsequent depressive phases. RNA sequencing was performed on iMG cells at each time point to identify differentially expressed genes related to mood state transitions.

**Results:** A thorough analysis of longitudinal gene expression data has led to the identification of three functional gene categories: “state-dependent genes”, “depression-to-mania transition genes (named: firing genes)”, and “mania-to-depression transition genes (named: extinguishing genes)”. A total of 168 firing, 59 extinguishing, and 77 state-dependent genes were identified. Notably, functional annotation revealed that, compared to the extinction gene set, the firing gene set was enriched in immune and inflammatory response pathways, particularly early-response cytokines such as IL1B and TNF.

**Conclusions:** Based on these findings, we propose that inflammatory immunomodulation by microglia contributes to mood switching in BD, especially in the process of depression-to-mania transition. The classification of genes by their relationship to state transitions offers a novel framework for understanding the molecular mechanisms underlying this complex disorder and may identify potential therapeutic targets to stabilize mood. Further validation with larger cohorts is warranted.

## Introduction

Bipolar disorder (BD) is characterized by two completely opposite symptoms, depression and mania. First described as “folie circulaire” by Jean-Pierre Falret in 1854, and concurrently as “folie à double forme” by Jules Baillarger^1,2^. Since the seminal reports by the two French psychiatrists 170 years ago, various biological research has been reported including pharmacological, genetic and brain imaging studies^2-5^. However, the pathophysiology underlying BD, especially in the context of switching mechanisms, is still not revealed^6-9^.

The inflammation hypothesis has been proposed as one of the pathological mechanisms, supported by some reports of activated microglia in BD^10-15^. We herein hypothesis that alterations in immunological conditions of microglia contribute to mania-to-depression and depression-to-mania transitions in BD. To address this hypothesis, given challenges associated with obtaining and analyzing live human microglia, we utilized a novel reverse-translational technique to generate directly induced microglia-like (iMG) cells, which are surrogate cells showing a microglial phenotype^16-22^. A pilot study using iMG cells from three patients with BD showed the significant expression differences of inflammation-related genes at manic and depressed states^23^. These previous findings have indicated that the gene expression profile of iMG cells can be altered by the states of BD. In the present study, iMG cells were repeatedly produced from a single patient with BD and a comprehensive gene expression analysis was conducted on the iMG cells to determine biomarkers associated with the switching of the states of BD.

## Materials and methods

This study received ethical approval from the Kyushu University ethics committee (25-84). The diagnosis of BD was determined by a psychiatrist and depressive symptoms were assessed with the Hamilton Rating Scale for Depression (HRSD)^24^ and manic symptoms with the Young Mania Rating Scale (YMRS)^25^. Total RNA was extracted from the iMG cells during each clinical phase and RNA sequencing was performed for comprehensive gene expression analysis. To evaluate the microglial function of iMG cells, we filtered previously reported microglia-related genes^26^ and differentially expressed genes (DEGs) were extracted as those with an absolute log_2_ fold change (|log□FC|) > 1. Because statistical testing was not feasible, genes were selected solely based on predefined fold-change criteria and are referred to as DEGs in this study. The detail methods are shown in the Supplementary Information.

## Results

### Case history

The case is a woman in her 40s with no significant developmental history or family history of psychiatric disorders. She presented with long-standing short sleep duration (<4 hours/night) and co-morbid obesity, hypertension, and dyslipidemia. Since her youth, she had experienced small mood swings, and in her mid-30s she began to have distinct depressive episodes accompanied by insomnia and anxiety. She also showed irritability and was unable to concentrate on her work, leading to her resignation. Concurrently, she received a diagnosis of BD and initiated a medical regimen, but her mood swings gradually worsened. During depressive phases, she exhibited withdrawal, decreased appetite, and suicidal ideation. Conversely, manic episodes were marked by euphoria and impulsive behavior. She reported seasonal fluctuations with worsening depression during winter. Throughout the approximately six-month study period, blood samples collected at three time points corresponded to a shift from depressive to manic state and then back to depression.

During the present study, her mood symptoms were tracked for approximately half a year and blood samples were collected at three separate points. The patient was treated with medication including valproic acid 600 mg, flunitrazepam 1 mg and suvorexant 20 mg during this study, and the patient’s states shifted from depressive to manic state, and then back to depressive state (Fig 1A).

**Figure 1.**
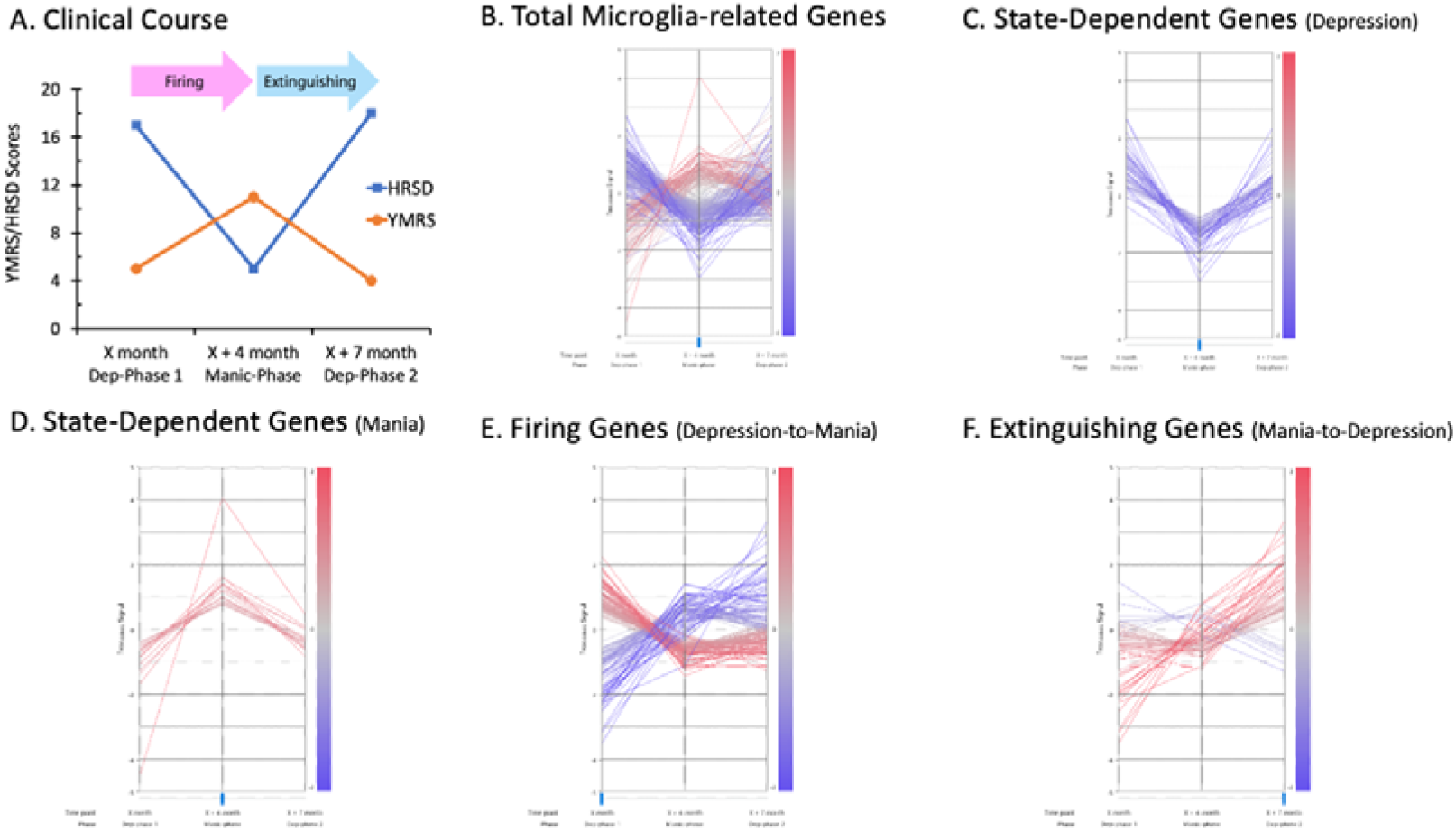
Longitudinal microglia-related transcriptional changes across recurrent depression–mania–depression transitions. [A] Clinical course and disease-phase transitions across the depression–mania–depression sequence. The depression-to-mania transition from Dep-Phase 1 to Manic-Phase was termed “Firing,” reflecting the patient’s subjective experience of becoming increasingly active, whereas the mania-to-depression transition from Manic-Phase to the subsequent Dep-Phase 2 was termed “Extinguishing.” [B] Microglia-related genes showing expression changes during at least one disease-phase transition. Centered expression profiles of all identified genes. [C] Depressive state-dependent genes. State-dependent genes showing higher expression during depressive phases than during the manic phase. [D] Manic state-dependent genes. State-dependent genes showing higher expression during the manic phase than during depressive phases. [E] Firing genes. Microglia-related genes showing expression changes during the “Firing (depression-to-mania)” transition after excluding state-dependent genes (C&D). [F] Extinguishing genes. Microglia-related genes showing expression changes during the “Extinguishing (mania-to-depression)” transition after excluding state-dependent genes (C&D).

### Gene expression analysis of the iMG cells

To characterize gene expression changes of iMG cells associated with mood transitions, the microglia-related DEGs (Fig. 1B) were categorized into three distinct functional groups. Genes exhibiting sustained differential expression across both depressive states and manic state were designated “depressive state-dependent genes” (Fig. 1C) (Fig. 1D). Following exclusion of state-dependent genes, the remaining genes showing differential expression during the depression-to-mania transition were designated “firing genes” (Fig. 1E), whereas those showing differential expression during the mania-to-depression transition were designated “extinguishing genes” (Fig. 1F).

The implementation of these criteria resulted in the identification of 168 firing genes, 59 extinguishing genes, 59 depressive state-dependent genes and 18 manic state-dependent genes (Table 1). Functional annotation of these microglia-related gene sets indicated representation of immune- and inflammatory-response–related processes, although the specific terms differed across categories.

**Table 1.**
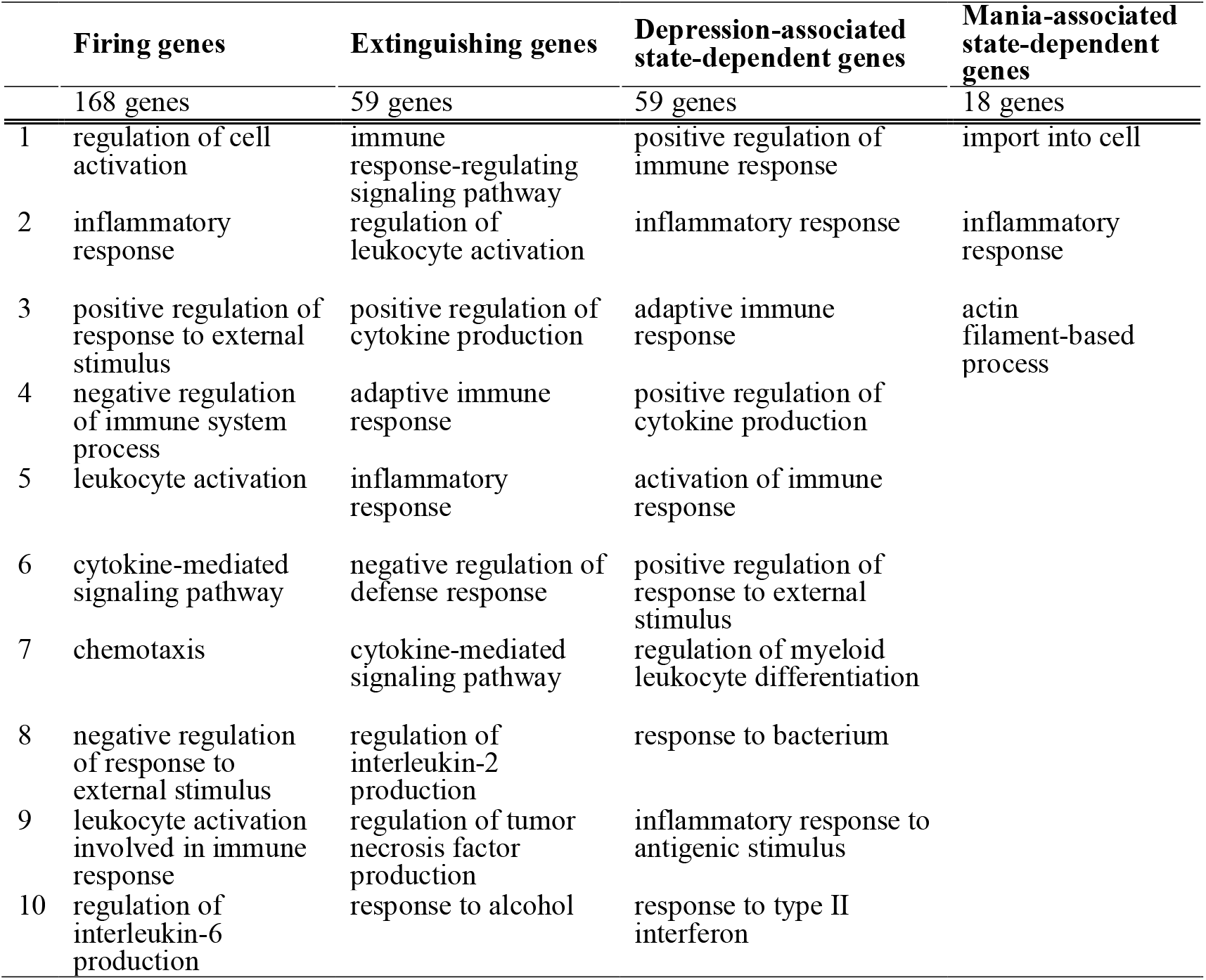
Top 10 GO Biological Process enrichment of microglia-related gene sets. GO Biological Process enrichment analysis was performed using Metascape for each microglia-related gene set (firing genes, extinguishing genes, and state-dependent genes defined across “Firing (depression-to-mania)” and “Extinguishing (mania-to-depression)” transitions). The table lists the top 10 GO terms for each gene set. Terms are ordered according to enrichment results generated by Metascape (over-representation analysis), without additional statistical interpretation in the present study. The manic state-dependent gene set yielded fewer than ten enriched GO terms, reflecting the limited size of the input gene list.

Firing genes were additionally represented in functional categories related to regulation of cell activation, whereas extinguishing genes were represented in categories related to immune response-regulating signaling pathway (Table 1). A comprehensive list of genes falling into each category is provided in the Supplementary Table. Notably, the firing genes category comprises early-response cytokines such as IL1B and TNF, indicating the presence of an inflammatory component that contributes to the transition towards mania.

## Discussion

This single case study using the iMG cells provides the first pilot evidence suggesting a microglia-derived immunomodulatory component to mood switching in BD. A key innovation of this work lies in our dissection of DEGs into three distinct categories: state-dependent, firing, and extinguishing genes. This approach has enabled more sophisticated understandings of transcriptional dynamics beyond simple correlations with mood state. The integration of longitudinal data from three distinct phases distinguished genes potentially involved in mood switches—the “firing” and “extinguishing” gene sets—from those which primarily reflect the current disease state.

Although the biological mechanisms underlying mood-state transitions remain poorly understood, accumulating evidence has implicated neuroimmune dysfunction and microglia-related alterations in the pathophysiology of BD. Previous studies have reported abnormalities in microglial markers, inflammatory signaling, and immune-related gene expression in patients with BD^12,23,27-31^. However, prior studies primarily compared established states, offering limited insight into transition-related molecular changes. Our findings demonstrate dynamic microglia-related gene expression accompanying recurrent mood shifts, with functional annotation revealing enrichment for immune and inflammation processes, suggesting a role for neuroimmunity in transcriptional regulation during transitions.

This study is limited by its single-patient design, which restricts generalizability. iMG cells represent an *in vitro* model and may not fully capture the complexity of resident human microglia within the central nervous system. The exploratory design and limited sampling points preclude definitive conclusions regarding causality or temporal directionality. Future studies with larger longitudinal cohorts, repeated sampling across multiple transitions, are needed to confirm these findings and distinguish transcriptional changes tracking clinical state from those driving it.

This pilot study has proposed a potential role for immunomodulation in driving mood switching in BD and highlight the potential of targeting inflammatory pathways. A subset of genes exhibited expression patterns corresponding to longitudinal changes in clinical symptom ratings,whereas other changes were not readily explained by differences between clinical states alone.

Although the biological significance of these findings remains uncertain, the present results demonstrate the feasibility of longitudinal transcriptomic analyses using patient-derived iMG cells and provide a framework for future studies investigating microglia-related transcriptional changes across mood-state transitions.

## Acknowledgments

This work was partially supported by Grant-in-Aid for Scientific Research: (1) The Japan Society for the Promotion of Science (KAKENHI; JP16H06279 (PAGS), JP18H04042, JP19K21591, JP20H01773, JP22H00494, and JP26H02458 to TAK), (2) The Japan Agency for Medical Research and Development (AMED; JP21wm0425010, and JP25wm0625322 to TAK) and (3) The Japan Science and Technology Agency CREST (JPMJCR22N5 to TAK). The funders had no role in the study design, data collection and analysis, decision to publish, or manuscript preparation.

