## Supplementary Information for "Transcriptomic signatures associated with mania-to-depression and depression-to-mania transitions in bipolar disorder: a case report using induced microglia-like (iMG) cells"

**Title:**

### Author information

Shogo Inamine, MS[1#], Sota Kyuragi, MD, PhD[1#], Masahiro Ohgidani, PhD[1&2], Tetsuaki Kimura, PhD[3], Ituro Inoue, PhD[3], Tomohiro Nakao, MD, PhD[1], Takahiro A. Kato, MD, PhD[4*]

### Patient

Recruitment for this study was conducted through the Mood Disorder/Hikikomori Clinic in the department of Neuropsychiatry at Kyushu University. This study was performed in line with the principles of the Declaration of Helsinki. Written informed consent was obtained from the patient included in the study. The ethics committee of Kyushu University approved the study (25-84). The participant has consented to the submission of the case report to the journal. The diagnosis of bipolar disorder was determined by a psychiatrist on the basis of the Diagnostic and Statistical Manual of Mental Disorders, Fourth Edition (DSM-IV). Depressive symptoms were assessed with the Hamilton Rating Scale for Depression (HRSD) and manic symptoms with the Young Mania Rating Scale (YMRS).

### Induction of iMG cells

Patient-derived induced microglia-like (iMG) cells were differentiated from peripheral blood monocytes as previously described[1]. Briefly, peripheral blood was collected in heparin-containing tubes. Peripheral blood mononuclear cells (PBMCs) were separated by density gradient centrifugation in Histopaque-1077 (Sigma Chemical Co., St. Louis, MO, USA). PBMCs were then transferred to RPMI-1640 (Nacalai Tesque, Kyoto, Japan) medium supplemented with 10% fetal bovine serum (Japan Bioserum Co Ltd., lot No. JBS-024963; heat-inactivated at 56°C for 30 min) and 1% Penicillin–Streptomycin (10,000 U/mL) (Thermo Fisher Scientific, Cat. No. 15140122). CD11b-positive cells were enriched by magnetic-activated cell sorting (MACS) using CD11b MicroBeads (Miltenyi Biotec, Bergisch Gladbach, Germany) according to the previously described procedure[1]. The isolated cells were seeded into 24-well culture plates at a density of $4 \times 10^5$ cells/mL (0.5 mL/well) and incubated overnight at 37°C in a humidified atmosphere containing 5% $CO_2$. After overnight incubation, the medium containing non-adherent cells was removed. Adherent cells (monocytes) were cultured in RPMI 1640 medium containing GlutaMAX (Thermo Fisher Scientific, Cat# 61870036), supplemented with 1% Antibiotic-Antimycotic (Thermo Fisher Scientific, Cat# 15240062), recombinant human GM-CSF (10 ng/mL; R&D Systems, Cat# 215-GM) and recombinant human IL-

34 (100 ng/mL; R&D Systems, Cat# 5265-IL) for 14 days to generate iMG cells.

**RNA sequencing**

RNA sequencing was performed to compare differences in gene expression patterns in each state of the patient. The total RNA was extracted from iMG cells using a High Pure RNA Isolation kit (Roche Diagnostics) according to the manufacturer's protocol. The sequencing libraries were prepared from 1 μg of total RNA with NEBNext Ultra Directional RNA Library Prep Kit for Illumina according to the manufacturer's instructions. Cluster amplification and 150 bp paired-end sequencing were performed according to the manufacturer's protocol (Illumina) for NovaSeq (Illumina, San Diego, California, USA).

**Data analysis**

All read data were checked with FastQC (v0.11.7) and trimmed using Trimmomatic (v-0.38). Transcript-level expression levels were quantified in the counts per million (CPM). The sequencing results were analyzed using Subio platform basic plug-in, version 1.24.5861 (Subio Inc., Kagoshima, Japan). Genes with CPM values below 3 in all samples were excluded. Values below the cutoff threshold were replaced with a fixed value of 2 and subsequently $\log_2$-transformed using the Log Transformation function in Subio Platform (base 2). To evaluate longitudinal transcriptional changes across disease phases, expression values were expressed as $\log_2$ fold changes relative to the manic-phase sample, which served as the reference sample. Genes showing an absolute $\log_2$ fold change ($|\log_2 FC|$) > 1 in either comparison (Dep-Phase 1 vs. Manic-Phase or Manic-Phase 2 vs. Dep-Phase) were retained for downstream analysis. Microglia-related genes were subsequently extracted using a previously reported human microglia-related gene set described by Gosselin et al. (2017)[2].

Genes were classified into four categories according to predefined longitudinal expression patterns across the depressive–manic–depressive sequence. Genes exhibiting sustained differential expression across both depressive states and manic state were designated "depressive state-dependent genes" (Fig. 1C) and "manic state-dependent genes". Following exclusion of state-dependent genes, the remaining genes showing differential expression during the depression-to-mania transition were designated "firing genes" (Fig. 1E), whereas those showing differential expression during the mania-to-depression transition were designated "extinguishing genes".

Over-representation analysis (ORA) was performed separately for each gene set using Metascape with the default background gene set. Gene Ontology Biological Process (GOBP) terms were retrieved, and the top 10 terms from the Metascape summary output were reported.

**Authors' contribution**

S.I and T.A.K. contributed to the study conception and design. S.K., T.N. and T.A.K. recruited the participant and/or collected clinical data. Material preparation, data collection and analysis were performed by S.I., S.K., M.O. and T.A.K.. T.K. and I.I. conducted the library preparation for RNA sequencing. The first draft of the manuscript was written by S.I. and S.K. and all authors commented on previous versions of the manuscript. All authors read and approved the final manuscript.

### Supplementary Table: Gene lists for each gene set.

| Firing genes | | | Extinguishing genes | Depressive state-dependent genes | Manic state-dependent genes |
|---|---|---|---|---|---|
| 168 genes | | | 59 genes | 59 genes | 18 genes |
| NFKBIZ | ATP7A | PCED1B | CD300LF | HK3 | B3GNT5 |
| BMF | ZNF846 | ACSM5 | CD300LB | IL1RN | GPR160 |
| CD300LB | C19orf38 | MERTK | CD4 | CYTH4 | TIFA |
| BTK | TNFAIP3 | TNFRSF10A | THEMIS2 | SPN | LY75 |
| RGS18 | MILR1 | TNFRSF10D | IL4I1 | SLC11A1 | MYO1G |
| IL4I1 | TBC1D2B | TNFRSF10C | GLUD1P3 | HLA-DRA | FOSB |
| C3 | CMKLR1 | TREML1 | GPR84 | HLA-DRB1 | XIRP1 |
| FPR1 | PLA2G7 | PYGL | DOK3 | KCTD12 | PLAG1 |
| TBX19 | EGR2 | KCNK13 | SERPINA1 | HLA-DRB5 | RGS1 |
| EMB | NAIP | VSIG4 | SLC37A2 | HLA-DPA1 | TRPC2 |
| RAB38 | CLEC5A | ALOX5 | HLA-DQB1 | ADCY7 | TVP23C |
| ADORA3 | MPEG1 | OTUD1 | HLA-DQA1 | TREM2 | USP53 |
| TRAF3IP3 | HAVCR1 | CLDN7 | PTGER4 | C1QA | HSPA6 |
| ENTPD1-AS1 | NCF1 | ALPK1 | OSM | CSF1R | SP140 |
| GPR65 | CCL4 | SORL1 | C1QB | MS4A7 | CXCL8 |
| GPR84 | CCL3 | SFXN2 | MRC2 | SELPLG | KCNJ5 |
| FUT4 | CD14 | IL1RAP | LILRB2 | MRC1L1 | IKBIP |
| SERPINB9 | CCR1 | PTPRJ | LILRB1 | HLA-DRB6 | IRAK2 |
| DPP4 | CD53 | PTPRC | P2RY6 | INPP5D | |
| PARVG | B4GALT1 | PIK3R5 | CEP135 | FKBP5 | |
| MLKL | KCNE3 | RNASE6 | TMC8 | SASH3 | |
| BLNK | TNFSF13B | ANKRD22 | SLC16A3 | FCGR1A | |
| MAF | PLAU | PIK3CG | CMTM7 | LAT2 | |
| DOCK2 | PLD4 | LINC01410 | CMKLR1 | C3AR1 | |
| TFCP2L1 | ELF4 | CD180 | CLEC5A | MS4A4A | |
| MYC | PLK3 | CD274 | MPEG1 | MNDA | |
| STARD5 | TYMP | PDCD1LG2 | CLEC2D | MRC1 | |
| PATL2 | EML4 | STEAP3 | CCL4 | CISH | |
| OSM | C1orf162 | CYTIP | CD14 | SLC9A7P1 | |
| IL12RB1 | PTCRA | BTBD19 | CARD11 | PIK3R6 | |
| DHRS9 | CEBPA | PLBD1 | ZMYND15 | ADAMTSL4 | |
| PLXDC2 | APOBR | LY86 | PTGS1 | C1QC | |
| FFAR4 | KCNQ1 | FGD2 | FCGR2C | SLA | |
| C5AR2 | RNF125 | CC2D2B | SIGLEC10 | AOAH | |
| IER3 | LCP1 | FRRS1 | LAIR1 | TLR8 | |
| ARHGAP15 | IGSF10 | NLRC5 | IFI30 | CIITA | |
| ARHGAP25 | RHBDF2 | CSF3R | CH25H | GPR34 | |
| MMP14 | SOWAHD | CHEK2 | LRRC25 | OLR1 | |
| RASGRP4 | PRAM1 | HPSE | PARP15 | TNFRSF11A | |
| TNF | PTGS1 | OLFML2B | ARHGAP4 | DEF6 | |
| LILRA2 | HSPA1B | STK26 | ACSM5 | ALOX5AP | |
| LILRB2 | RASAL3 | SLCO2B1 | GIMAP8 | MS4A14 | |
| LILRB4 | LPCAT2 | NCKAP1L | VSIG4 | FOLR2 | |
| LILRB1 | CSF2RB | ITGB3 | ALOX5 | ATP8B4 | |
| DZIP1L | FCGR3A | TGFBR2 | MYO7A | ST8SIA4 | |
| PDE3B | FCGR1CP | A2M | CTSC | SIGLEC14 | |
| RCN3 | ABHD15 | SIRPB1 | SIGLEC7 | PLA2G4A | |
| IFNLR1 | SIGLEC10 | SSH2 | KCNMB1 | GSN-AS1 | |
| IL1B | LINC00921 | ABCA1 | ZC3H12A | M1AP | |
| OXER1 | LNPEP | ST14 | RNASET2 | SUSD3 | |
| TLR2 | JAK3 | | ITGB2-AS1 | TMEM52B | |
| TLR1 | IFI16 | | N4BP2 | HSPA7 | |
| IL15RA | LRRC25 | | FBP1 | SCIMP | |
| CCL3L3 | RUBCNL | | FGL2 | STAB1 | |
| ITPRIPL1 | LIPG | | DDX60L | LINC01160 | |
| LPAR6 | RAB39A | | NLRP3 | CYSLTR1 | |
| PAG1 | GPRIN3 | | CARD9 | P2RY13 | |
| RGL3 | CD300A | | CASS4 | MS4A6A | |
| RGS2 | CPVL | | CASP1 | FCGBP | |
